# The *Escherichia coli* Radical SAM Enzyme YhcC Replaces MnmC in mnm^5^s^2^U tRNA Modification During Anaerobic Growth

**DOI:** 10.64898/2026.05.18.725915

**Authors:** Kaleb Boswinkle, Paige N. Roehling, Colbie Reed, Jasmine Narakorn, Thomas Carell, Agnieszka Dziergowska, Patricia C. Dos Santos, Jeffrey S. Mugridge, Valérie de Crécy-Lagard

**Author notes:** These authors contributed equally (Kaleb Boswinkle, Paige N. Roehling).

## Abstract

tRNA wobble uridines are heavily modified to influence anticodon-codon pairing and tune anticodon stem-loop structure for efficient, accurate translation. Many bacteria and some archaea modify wobble uridines with either a 5-carboxymethylaminomethyl (cmnm^5^) or 5-methylaminomethyl (mnm^5^) group, often together with a 2-thio (s^2^) moiety. Bacteria utilize the conserved MnmEG complex to produce cmnm^5^U, which is further converted to mnm^5^U by non-orthologous enzymes in different lineages. *Escherichia coli* uses the bifunctional enzyme MnmC to demodify cmnm^5^U to nm^5^U and subsequently methylate nm^5^U to mnm^5^U, whereas *Bacillus subtilis r*elies on the radical SAM (rSAM) enzyme MnmL and the stand-alone methylase MnmM. Although *E. coli* and related bacteria encode homologs of MnmL, the function of the *E. coli* homolog, YhcC, remained unknown. Here, we show that YhcC is required for cmnm^5^s^2^U demodification in vivo during anaerobic growth, whereas the equivalent MnmC-dependent reaction requires O_2_ and occurs only aerobically. In vitro, purified [4Fe-4S]-reconstituted YhcC binds tRNA and catalyzes nm^5^s^2^U-tRNA synthesis from cmnm^5^s^2^U-tRNA. Together, these results define the previously unknown function of the *E. coli* rSAM enzyme YhcC and demonstrate that it replaces MnmC under anaerobic conditions to generate nm^5^s^2^U. These parallel pathways reveal how *E. coli* maintains synthesis of a critical wobble-base modification under both aerobic and anaerobic growth conditions.

**General audience summary:** Transfer RNA (tRNA) molecules serve as critical components in protein synthesis through their direct interaction with the ribosome and messenger RNA (mRNA). During protein synthesis, tRNA molecules utilize an anticodon composed of three bases that recognize a complementary mRNA codon. At the 3’CCA end of tRNA, the amino acid corresponding to the anticodon sequence is incorporated into the growing polypeptide chain. While canonical Watson-Crick base pairing, pairing between U:A and G:C, occurs between the second and third bases of the anticodon and the corresponding bases of the codon, there is increased flexibility in the ribosome at the first position of the anticodon. This results in non-canonical base pairing and allows a single tRNA molecule to recognize multiple codons. Modification at this site restricts or enhances this “wobble” base pairing. Bacteria often incorporate mnm^5^s^2^U34 into various tRNAs to facilitate proper and efficient translation. In *E. coli*, three proteins are responsible for this hypermodification pathway: MnmE and MnmG convert s^2^U to cmnm^5^s^2^U, while MnmC subsequently removes the carboxymethyl group to generate nm^5^s^2^U and methylates this intermediate into the ultimate product, mnm^5^s^2^U. We found that an additional enzyme, YhcC, is required for mnm^5^s^2^U synthesis in anaerobic conditions, specifically at the cmnm^5^s^2^U demodification step. Previous studies showed that MnmC-catalyzed demodification is dependent on FAD for the initial oxidation of tRNA substrate, generating FADH_2_. We propose that FADH_2_ recycling is dependent on O_2_ to regenerate FAD, allowing multiple catalytic turnovers. We show that, in vitro, O_2_ strongly promotes cmnm^5^s^2^U demodification by MnmC, supporting a new model of mnm^5^s^2^U synthesis with alternative aerobic and anaerobic routes. Conservation of multiple enzymes that perform similar chemistry, albeit under differing environmental conditions, highlights the importance of maintaining wobble base modifications as well as the ability of bacteria to adapt to their surroundings.

## Introduction

Transfer RNAs (tRNAs) are essential for protein synthesis across all domains of life, bridging messenger RNA (mRNA) codons with their corresponding amino acids. To ensure proper folding, interaction with protein partners, and accurate decoding during translation, tRNAs undergo extensive post-transcriptional modification (1, 2). In particular, position 34, known as the wobble base, which pairs with the third nucleotide of the codon, is heavily modified with diverse chemical nucleobase modifications (3, 4). These wobble base modifications modulate decoding efficiency as well as accuracy by expanding or restricting codon-anticodon base pairing in the ribosome (5). This is especially important for decoding of two-codon family boxes, where the identity of the last base of the codon as either a pyrimidine or purine dictates the corresponding amino acid. Wobble base uridines (U34) specifically are nearly invariably modified across all organisms, in part due to the inherent base pairing flexibility of this base (6). Most wobble uridine modifications are derivatives of 5-methyluridine (xm^5^) and also often contain a thio group at the C2 position (s^2^). In bacteria such as *E. coli* and *B. subtilis*, tRNAs that decode NNA/NNG codons typically contain a wobble uridine with either a 5-carboxymethylaminomethyl (cmnm^5^) or a 5-methylaminomethyl (mnm^5^) group, in addition to a 2-thio (s^2^) modification (7, 8). RNA modifications are important for various stress responses and adaptations. Recent studies have shown that *mnmE* mutants lacking cmnm^5^ exhibit a strong growth defect under acid stress as well as decreased cell motility (9). The importance of wobble uridine modifications is further highlighted by synthetic lethality in both *E. coli* and *S. enterica* LT2 when both *mnmE* and *mnmA* are deleted, resulting in completely unmodified U34 in various tRNAs. (10, 11).

The biosynthesis of (c)mnm^5^(s^2^)U34 modifications in bacteria begins with the MnmEG complex, which utilizes methylenetetrahydrofolate and either glycine (‘glycine route’) or ammonium (‘ammonium route’) to install a cmnm^5^ or nm^5^ (5-aminomethyluridine) moiety, respectively (12, 13) (**Fig. 1**). While MnmEG activity with ammonium was first shown *in vitro*, it was also found that MnmEG seems to prefer this activity over the glycine route in stationary phase *in vivo* (13, 14). In contrast, the glycine route was preferred in exponential phase (14). In *E. coli*, a single bifunctional enzyme, MnmC, then converts cmnm^5^(s^2^)U to nm^5^(s^2^)U and finally to mnm^5^(s^2^)U (15). The C-terminal FAD-dependent MnmC1 oxidoreductase domain transforms cmnm^5^(s^2^)U to nm^5^(s^2^)U and subsequently the N-terminal SAM-dependent MnmC2 methyltransferase domain methylates nm^5^(s^2^)U to mnm^5^(s^2^)U (16, 17). However, bifunctional MnmC is mainly restricted to the Pseudomonadota, whereas Gram-positive bacteria, such as *Bacillus subtilis*, instead rely on the recently discovered MnmL and MnmM enzymes to perform similar reactions (18–20). MnmL is a member of the radical SAM (rSAM) enzyme superfamily, a large and versatile group of enzymes that catalyze difficult chemical transformations often involving *sp^3^* or *sp^2^*hybridized carbon centers via radical intermediates (21, 22). Interestingly, *E. coli* and related bacteria that encode MnmC also encode a homolog of MnmL, which is named YhcC in *E. coli*. As the deletion of *mnmC* in *E. coli* results in cmnm^5^(s^2^)U accumulation with no nm^5^(s^2^)U production under standard growth conditions, the function of these MnmL homologs in *E. coli* and related bacteria remained unclear (19).

**Figure 1.**
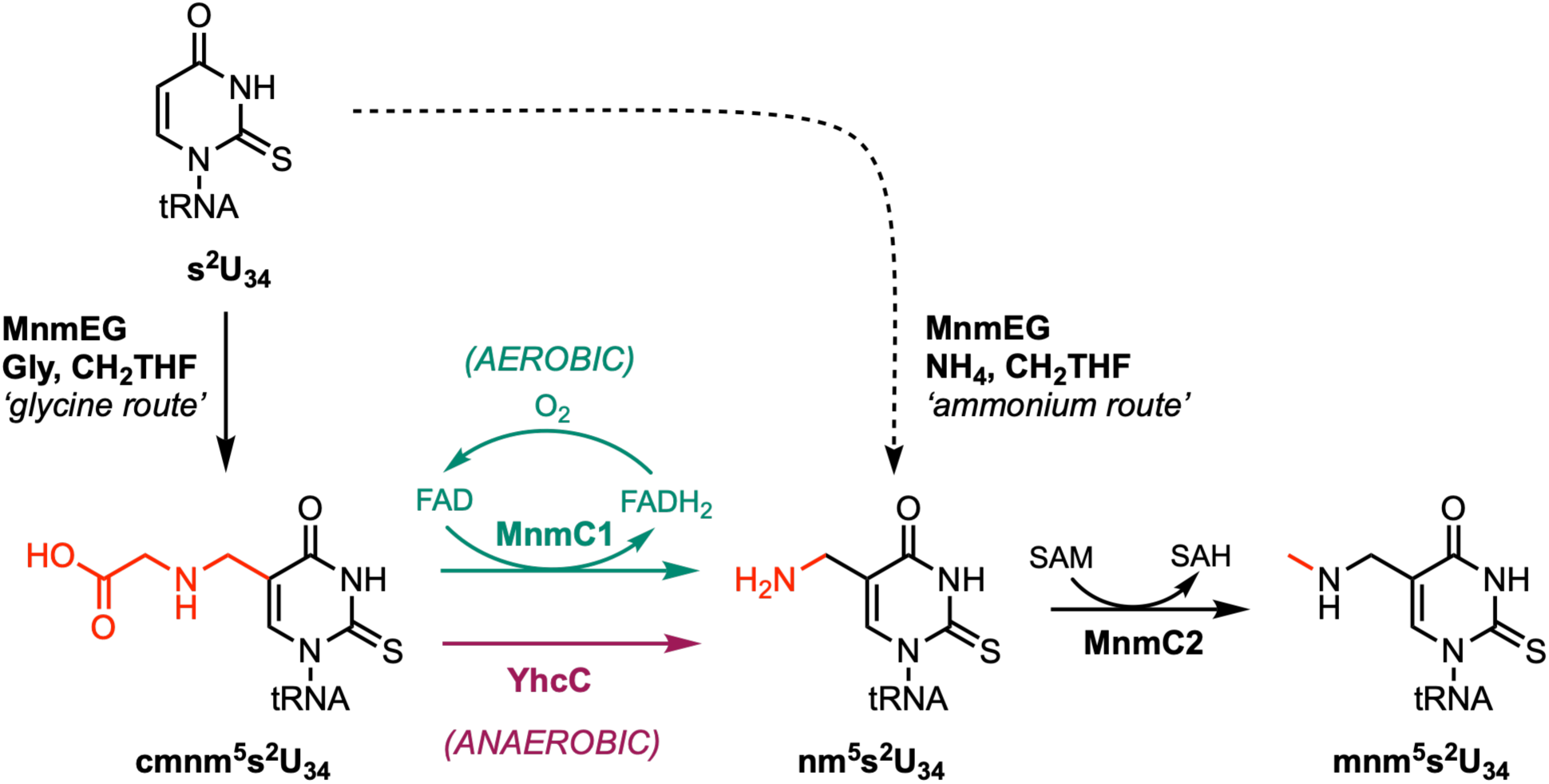
An updated overview of the mnm^5^s^2^U biosynthesis pathway in *E. coli*. MnmEG catalyzes one of two reactions: the ‘glycine route’ forms cmnm^5^s^2^U34 from U34 with glycine, tetrahydrofolate, and additional cofactors GTP, FAD, and NADH, and the postulated ‘ammonium route’ directly forms nm^5^s^2^U34 with ammonium in place of glycine. However, our results indicate that the previously reported stationary-phase accumulation of nm^5^s^2^U in *E. coli* can be explained by YhcC activity rather than requiring invocation of an ammonium-route mechanism (dotted line). MnmC contains two active sites: the MnmC1 active site is an FAD-dependent oxidoreductase that demodifies cmnm^5^s^2^U34 to nm^5^s^2^U34. We show that the MnmC1-catalyzed formation of nm^5^s^2^U is oxygen-dependent and only occurs under aerobic conditions (teal), and that the rSAM enzyme MnmL (formerly named YhcC) replaces MnmC1 under anaerobic growth conditions (purple). Finally, the MnmC2 methyltransferase domain of MnmC methylates nm^5^s^2^U34 to form mnm^5^s^2^U34 under both aerobic and anaerobic conditions.

Here, we determine the function of the previously uncharacterized *E. coli* rSAM enzyme YhcC by a combination of bioinformatics, genetics, and biochemistry. We show that YhcC is required for the conversion of cmnm^5^s^2^U to nm^5^s^2^U under anaerobic growth conditions in *E. coli*, and that the analogous reaction carried out by MnmC1 is likely inactive under anaerobiosis due to an O_2_ requirement for FADH_2_ oxidation during the FAD-dependent MnmC1 catalytic cycle. Additionally, we investigate the biochemistry of YhcC and show *in vitro* tRNA binding and conversion of cmnm^5^s^2^U_34_ to nm^5^s^2^U. Finally, we revisit the postulated growth-phase dependent activities of MnmEG in light of the presence of an anaerobically-regulated, YhcC-dependent route for nm^5^s^2^U synthesis and show that YhcC is in fact responsible for the reported ‘ammonium route’ activity of MnmEG during stationary phase. These studies redefine the tRNA wobble base mnm^5^s^2^U modification biosynthesis and its regulation under aerobic and anaerobic growth in *E. coli*.

## Results

### Gene neighborhood analysis, transcriptional regulation, and RNA-Seq data link YhcC to anaerobiosis

To gather functional knowledge on the *E. coli yhcC* gene, we surveyed the literature and available transcriptomic data. It was reported that *yhcC* is positively regulated by the FNR (fumarate and nitrate reductase regulation) transcription factor in anaerobic growth by ChIP-Seq and RNA-Seq experiments (23, 24). FNR regulates the switch from aerobic to anaerobic metabolism in *E. coli* and related facultative anaerobic bacteria and is required for the expression of genes involved in anaerobic respiration (25, 26). Besides those involved in anaerobic respiration, additional genes are either activated or repressed by FNR and a total of over 100 genes are regulated by FNR in *E. coli*. YhcC binds a [4Fe-4S] cluster and cleaves SAM to methionine and 5’-deoxyadenosine in the presence of a chemical reductant, confirming that it is a rSAM enzyme (27). However, the reaction catalyzed by the enzyme remained unknown.

Genes involved in similar processes are often co-localized on bacterial chromosomes (28). Analysis of the gene neighborhood of *yhcC* in *E. coli* and other gammaproteobacteria shows that *yhcC* is often found adjacent to *arcB*, which encodes the sensor protein of the two-component Arc (aerobic respiration control) system (**Fig. S1**). The Arc system mainly represses aerobic respiration genes under anaerobic conditions (29). This association is in contrast to *mnmL* found in Gram positive bacteria, which is often adjacent to or fused with *mnmM* (19). Collectively, these analyses suggest YhcC may be active under anaerobic metabolic conditions as opposed to aerobic conditions.

### Sequence and structural analysis do not differentiate *E. coli* YhcC from *B. subtilis* MnmL

We gathered YhcC/MnmL sequences (IPR005911) from organisms that, like *E. coli*, encode *mnmC* and from organisms that do not, like *B. subtilis,* and constructed both a small and large multiple sequence alignment. The former allowed us to search for motifs that could separate proteins found in organisms that either lack or encode MnmC proteins. We failed to identify any residues that could differentiate the two groups (**Fig. S2**). We then used AlphaFold3 to predict the structure of *E. coli* YhcC and *B. subtilis* MnmL and the backbones aligned with an RMSD of 3.84 angstroms (**Fig. S3B**) (30). Subsequently, we used the large multiple sequence alignment to identify conserved residues. This enabled us to locate the approximate location of the YhcC active site based on the presence of highly conserved cysteine residues, which are used by rSAM enzymes to coordinate a [4Fe-4S] cluster. We found that the predicted structure of EcYhcC contains a cleft lined with positively charged residues adjacent to the potential active site and hypothesized that these residues may be involved in nucleic acid binding (**Fig. S3**). Two of these residues, K21 and R279 are highly conserved and are located at the opening of this cleft (**Fig. S3**).

Given the apparent association between *yhcC* and anaerobic metabolism and its close resemblance to recently experimentally validated MnmL enzymes, we hypothesized that YhcC in *E. coli* and related organisms may be a true MnmL ortholog that is active under anaerobiosis. Since previous work on *E. coli ΔmnmC* strains in relation to the biosynthesis of tRNA modifications only analyzed cells grown aerobically, the role of *yhcC* may have been overlooked.

### *E. coli* YhcC is essential for mnm^5^s^2^U synthesis under anaerobic conditions

Previous studies had shown that deletion of *mnmC* in *E. coli* results in the absence of mnm^5^s^2^U and the accumulation of cmnm^5^s^2^U in tRNA during exponential phase, or the accumulation of nm^5^s^2^U in tRNAs extracted during stationary phase (14, 19). We confirmed these observations by analyzing modification profiles using LC-MS on bulk tRNAs extracted from *E. coli* cells grown aerobically to either exponential phase (OD_600_ ≈ 0.6) (**Fig. 2A**) or stationary phase (OD_600_ ≈ 3.0) (**Fig. 2C**). Furthermore, we showed that deletion of *yhcC* does not affect the ability of *E. coli* to produce mnm^5^s^2^U in tRNAs extracted from cells in both exponential and stationary phases under aerobic conditions (**Fig. 2A and 2C**).

**Figure 2.**
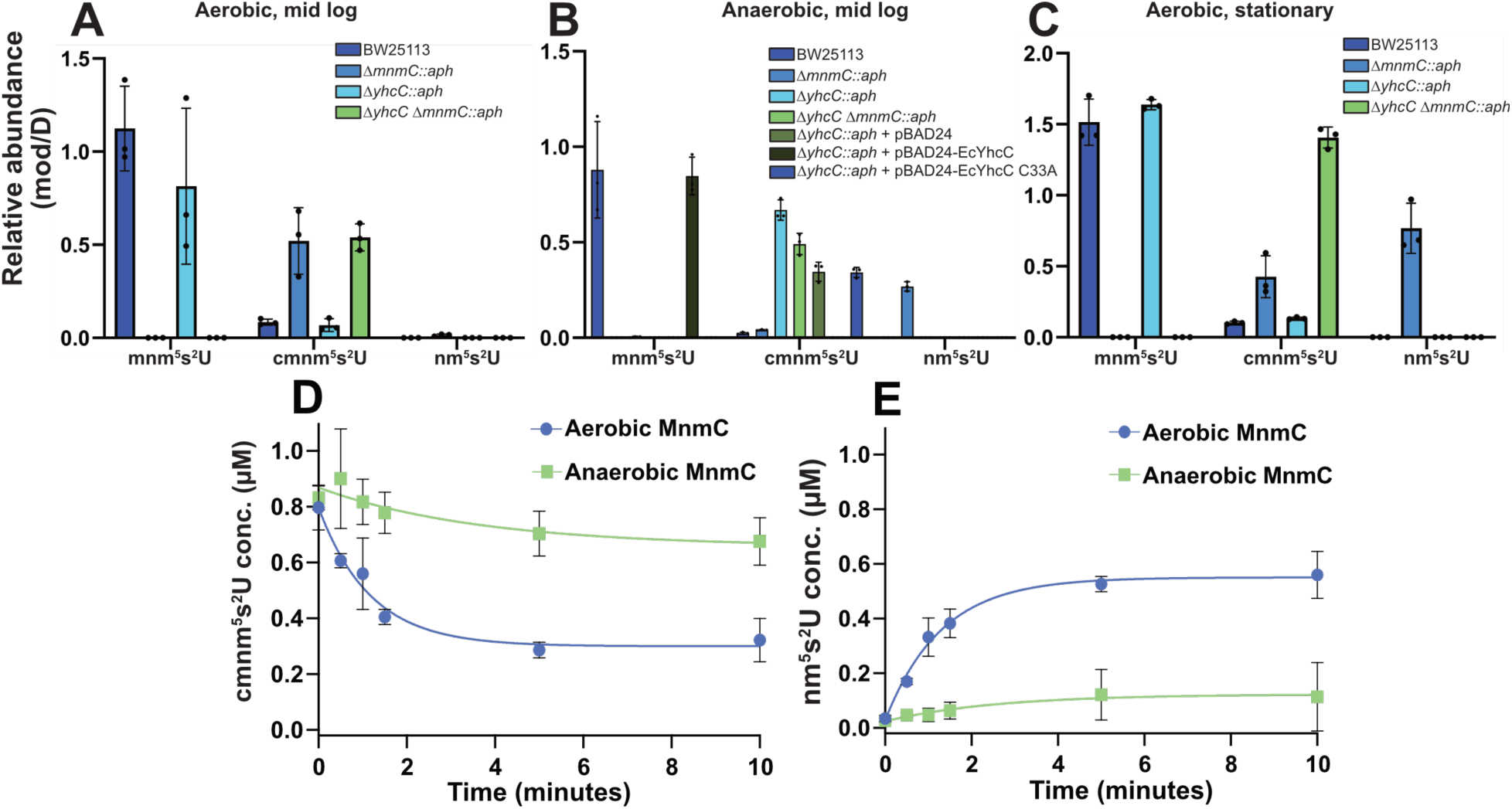
Accumulation of mnm^5^s^2^U tRNA intermediates under aerobic and anaerobic conditions. **A-C** (top): cmnm^5^s^2^U, nm^5^s^2^U, and mnm^5^s^2^U abundances, normalized by the amount of dihydrouridine (D) in the same sample, in tRNAs purified from *E. coli* in triplicate cultures at different stages of growth under either aerobic or anaerobic conditions. Bars represent mean values and error bars represent the standard deviation. **A**: *E. coli* grown aerobically and harvested at mid-log phase, **B**: *E. coli* grown anaerobically and harvested at mid-log phase, and **C**: *E. coli* grown aerobically and harvested at stationary phase. **D** and **E**: *In vitro* MnmC activity assays under aerobic and anaerobic conditions, monitoring disappearance of substrate cmnm^5^s^2^U (**D**) and appearance of product nm^5^s^2^U (**E**) in purified tRNA pools obtained from Δ*mnmC E. coli* cells with recombinant MnmC enzyme. Data are shown as mean values ± SD and are fitted to a single-phase exponential equation.

Next, we tested the hypothesis that *yhcC* is required for cmnm^5^s^2^U to nm^5^s^2^U conversion under anaerobic conditions by repeating the above analysis with cells grown under anoxic conditions (**Fig. 2B**). We found that deleting *mnmC* or *yhcC* led to the absence of mnm^5^s^2^U in bulk tRNA. However, nm^5^s^2^U accumulated in the Δ*mnmC* strain but not in the Δ*yhcC* strain under anaerobic growth. We did not detect the nm^5^s^2^U intermediate when both *mnmC* and *yhcC* were deleted (*ΔmnmC/yhcC*). Additionally, cmnm^5^s^2^U accumulated in tRNAs extracted from the Δ*yhcC* mutant grown under anoxic conditions despite the presence of *mnmC*. To confirm that the loss of mnm^5^s^2^U in anaerobically-grown *E. coli ΔyhcC* was not due to a polar effect, we complemented the loss of *yhcC* by expressing the wild-type *yhcC* gene using pBAD24 (**Fig. 2B**). The complemented *E. coli ΔyhcC* strain produced only mnm^5^s^2^U, in contrast to both the knockout and the wild-type strain, which had detectable levels of cmnm^5^s^2^U. Finally, we mutated one of the cysteine residues (C33) likely involved in coordinating the rSAM [4Fe-4S] cluster to alanine. Expression of this variant did not recover the *ΔyhcC* phenotype, as it accumulated only cmnm^5^s^2^U, showing that the conversion of cmnm^5^s^2^U to nm^5^s^2^U relies on YhcC catalysis.

The apparent lack of YhcC activity during aerobic growth could either be due to a lack of *yhcC* transcription, or possibly due to the instability of the iron-sulfur cluster in the presence of molecular oxygen. To investigate these two scenarios, we expressed *yhcC* in pBAD24 containing the arabinose-inducible P*_bad_* promoter under aerobic conditions in the double *ΔmnmC*/*ΔyhcC* background. We also overexpressed *mnmL* from *Bacillus subtilis*, which has previously been reported to be active when expressed in *E. coli* under aerobic conditions (19). We found that while the strains overexpressing either *yhcC* or *mnmL* had decreased levels of cmnm^5^s^2^U relative to the wild-type strain transformed with the empty vector, we surprisingly could not detect nm^5^s^2^U or mnm^5^s^2^U in either strain (**Fig. S4**). However, the strain overexpressing the C33A *yhcC* mutant reversed the observed decreased cmnm^5^s^2^U level chemotype, and instead, the tRNA modification profile of this strain appeared closer to the wild-type control. While we were unable to detect nm^5^s^2^U in the *yhcC*-overexpression strain, this supports the idea that YhcC is likely active under aerobic conditions. While the fate of the depleted substrate remains unresolved, possibilities are that the nm^5^s^2^U-containing tRNA is unstable in this growth condition, or nm^5^s^2^U could be converted to a modified nucleoside that we were unable to identify. We also were unable to detect the non-thiolated versions of the modified nucleosides.

Our epitranscriptomic tRNA analysis replicated an earlier finding that nm^5^s^2^U accumulates in tRNAs extracted from a Δ*mnmC* mutant grown to stationary phase (**Fig. 2C**) (14). However, we did not detect this modification is in the double mutant, Δ*mnmC*/*yhcC,* and instead found an increase of the cmnm^5^s^2^U precursor. As stationary phase is the only growth condition in which the ammonium route of MnmEG has been proposed to be active, our findings challenge the idea that the nm^5^ moiety can be inserted directly by MnmEG *in vivo* (14). These results demonstrated that the formation of mnm^5^s^2^U in the absence of oxygen requires both the rSAM enzyme YhcC for the synthesis of the nm^5^s^2^U intermediate and the methyltransferase activity of MnmC2.

The FAD-dependent activity of MnmC1 was detected only under conditions where oxygen is present. We hypothesized this could be due to a requirement for O_2_ to recycle the FADH_2_ cofactor. Since previously reported *E. coli* MnmC activity assays had not tested the oxygen dependence of the MnmC1 oxidoreductase, we tested the O_2_ requirement of the MnmC1-catalyzed cmnm^5^s^2^U to nm^5^s^2^U conversion (**Fig. 2**) (17). Using hypomodified tRNA isolated from *E. coli* Δ*mnmC* overexpressing tRNA^Glu^_UUC_, which accumulates cmnm^5^s^2^U, we carried out *in vitro* activity assays with recombinant MnmC under both aerobic and anaerobic (< 1 ppm O_2_) conditions, monitoring conversion of substrate cmnm^5^s^2^U into product nm^5^s^2^U with UHPLC-MS. Consistent with our *in vivo* data, we find that MnmC1 activity is significantly reduced under anaerobic conditions, suggesting that O_2_ is indeed required for FADH_2_ recycling and robust cmnm^5^s^2^U to nm^5^s^2^U tRNA modification activity by MnmC.

### *E. coli* YhcC *in vitro* tRNA binding and catalytic activity

As our genetic studies in *E. coli* showed a clear role for *Ec*YhcC in nm^5^s^2^U-tRNA synthesis, we next set out to show directly that YhcC can bind tRNA and catalyze the conversion of cmnm^5^s^2^U to nm^5^s^2^U *in vitro*. For these experiments, we first overexpressed and purified *Ec*YhcC from *E. coli* cells and chemically reconstituted its [4Fe-4S] cluster under anaerobic conditions (**Fig. S5**). To measure YhcC-tRNA interactions, we used electrophoretic mobility shift assays (EMSAs) with purified YhcC and unmodified *E. coli* tRNA^Lys^_UUU_ prepared by *in vitro* transcription and found that YhcC binds tRNA with a *K*_D_ of approximately 5 μM (**Fig. S6B**). Next, we carried out tRNA modification assays using a substrate cmnm^5^s^2^U34-enriched tRNA pool and our purified, reconstituted YhcC under anaerobic conditions, monitoring cmnm^5^s^2^U to nm^5^s^2^U conversion by UHPLC-MS (**Fig. 3**). These reactions showed a clear YhcC- and SAM-dependent consumption of substrate cmnm^5^s^2^U over time, and the concomitant production of product nm^5^s^2^U, providing direct *in vitro* evidence that YhcC catalyzes the cmnm^5^s^2^U to nm^5^s^2^U reaction on tRNA.

**Figure 3.**
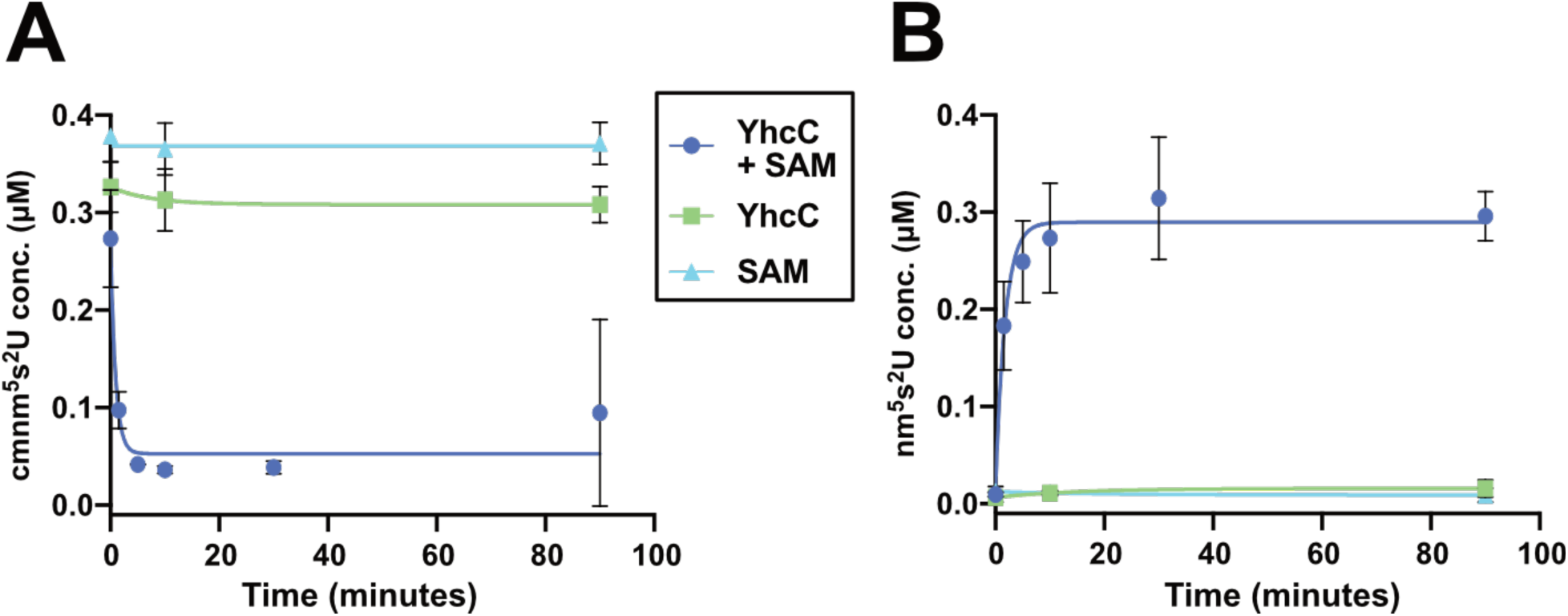
YhcC *in vitro* assays catalyzing the conversion of cmnm^5^s^2^U to nm^5^s^2^U tRNA. Activity assays were performed under anaerobic conditions with purified, reconstituted YhcC (1 μM) and cmnm^5^s^2^U-enriched tRNA substrate (3.5 μM total tRNA concentration) obtained from *E. coli* Δ*mnmC* cells overexpressing tRNA^Glu^. Reactions were quenched at different times with EDTA and tRNA is digested to single nucleosides for UHPLC-MS analysis. YhcC catalyzes the disappearance of substrate cmnm^5^s^2^U modifications (**A**) and production of nm^5^s^2^U modifications (**B**) in an enzyme- and SAM-dependent manner. Data are shown as mean values ± SD and are fit to a single-phase exponential equation.

### Phylogenetic analysis of the occurrence of MnmL and MnmC homologs and their association with physiological oxygen utilization status

After finding that YhcC is required in *E. coli* for the oxidative demodification of cmnm^5^s^2^U specifically under anaerobic conditions we were interested to investigate if the dual use of YhcC (MnmL) and MnmC is a general strategy utilized by facultative bacteria. We surveyed 1,322 representative bacterial genomes spanning 54 phylogenetic groups for the presence of MnmC and/or MnmL homologs utilizing the KEGG and COG databases (31, 32). We found that 27.2% encode MnmL, 18.2% encode MnmC, 4.4% encode both and the remaining 59% do not encode either of these genes. While six phyla were found to encode MnmC homologs, the vast majority of these were found to be present in Pseudomonadota (232 of 240 homologs) (**Fig. S7**). In contrast, MnmL homologs are more common and found in a much more phylogenetically diverse set of bacteria (**Fig. S8**). Next, we used the BactoTrait and Bac*Dive* databases to curate physiological oxygen utilization data and recovered data for 707 of 1,322 organisms (**Fig. S9**) (33, 34). Organisms were determined to fall into one of four classes based on MnmL and MnmC incidence: “neither”, “MnmL-only”, “MnmC-only”, or “both”. Leveraging the KEGG Orthology (KO) database, we determined that MnmL-only organisms tended towards anaerobic bacteria, while MnmC-only organisms as well as organisms not possessing either MnmL or MnmC were predominantly aerobic (**Fig. 4A**). Those in which MnmC and MnmL co-occurred were found to trend towards a facultatively anaerobic average. Additionally, the incidence of MnmL strongly decreased with increasing oxygen requirements of respective genomes, while the incidence of MnmC did not exhibit a strong correlation with oxygen requirement status (**Fig. 4B-C**). These findings support the possibility that facultative organisms may more generally alternate between YhcC and MnmC. However, this is not a widely utilized strategy within bacteria, as the co-occurrence of MnmC and MnmL is restricted mainly to Pseudomonadota bacteria.

**Figure 4.**
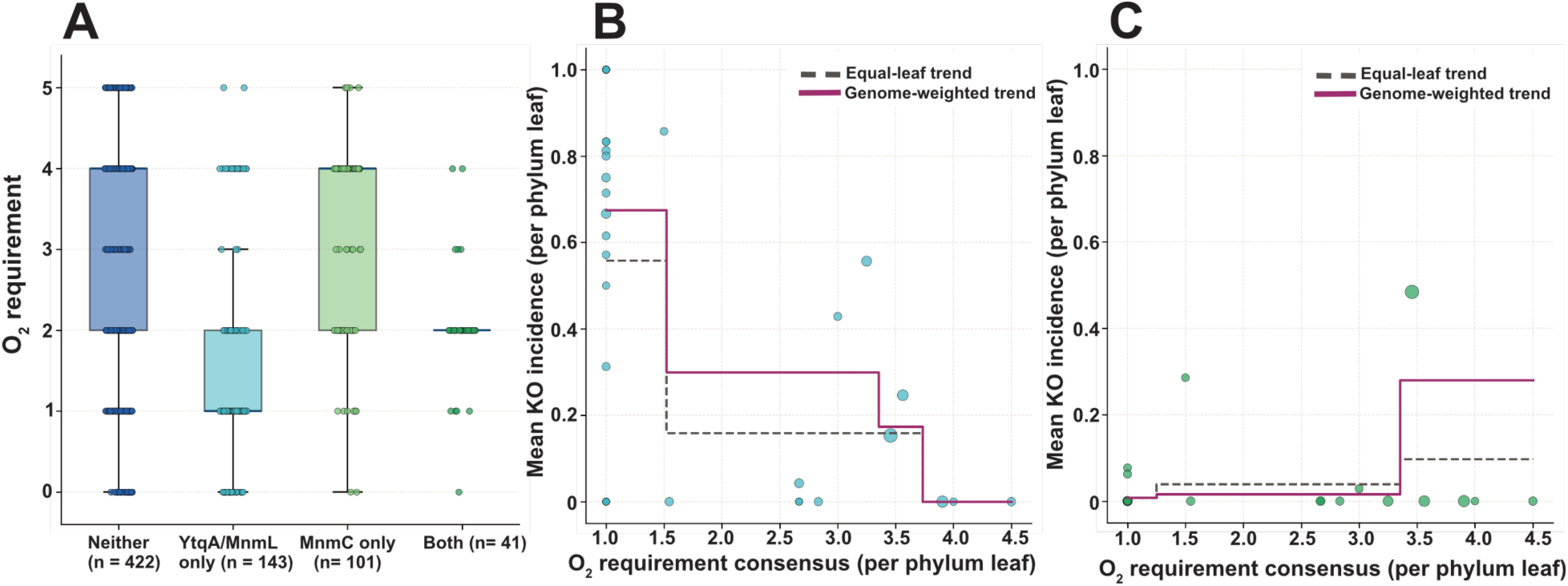
MnmL (K07139) and MnmC (K15461) incidence vs. oxygen requirement. **A**: Box and whisker plots representing per-genome oxygen score by MnmC and MnmL incidence. Boxes represent interquartile ranges (IQR) and whiskers represent values within 1.5 × IQR. Panels **B** and **C**: summary-leaf (phylum) mean KO incidence vs. oxygen consensus, for K07139 (**B**) and K15461 (**C**) respectively, each with a linear fit. Point size is monotonically scaled by genome count; square-root scaling compresses the large differences between phyla (minimum=1, area= 24 points; median= 5, area= ∼29 points; Pseudomonadota=262, area= ∼85 points). Genome-weighted trendline is shown in magenta.

## Discussion

Previous studies had implicated MnmL (previously called YtqA) from *Bacillus subtilis*, *Streptococcus mutans*, and *Streptococcus pneumoniae*, in mnm^5^s^2^U synthesis (19). However, while genetic studies show that MnmL from Bacillota is required for the synthesis of nm^5^s^2^U from cmnm^5^s^2^U, this had not been confirmed biochemically (19). We show here that the *yhcC* gene in *E. coli* encodes an *mnmL* ortholog that is required for nm^5^s^2^U formation specifically under anaerobic conditions, even in the presence of *mnmC*. Therefore, *yhcC* should be renamed to *mnmL* in *E. coli* and in related facultative anaerobes. Indeed, both our *in vivo* and *in vitro* data show that the FAD-dependent oxidoreductase (or MnmC1) activity of MnmC is not present under anaerobic conditions. Notably, the crystal structure of MnmC revealed that the MnmC1 active site is highly similar to that of *Bacillus subtilis* glycine oxidase, which utilizes molecular oxygen to oxidize the reduced flavin cofactor (35, 36). In the absence of oxygen, assays with glycine oxidase from *Marinomonas mediterranae* with several alternative electron acceptors failed to detect activity (37). However, facultative anaerobes must adapt to conditions lacking oxygen, and utilize anaerobic metabolism. Some FAD-dependent oxidases can utilize alternative electron acceptors, such as fumarate in the case of L-aspartate oxidase (38). However, the lack of MnmC1-dependent cmnm^5^s^2^U demodification activity in the *ΔyhcC* strain under anaerobic conditions suggests a stringent dependency on O_2_ for the oxidoreductase activity of MnmC. Indeed, *in vitro* kinetic measurements of MnmC in the presence or absence of oxygen show that O_2_ greatly enhances activity. Interestingly, despite the lack of MnmC1-dependent activity under anaerobic conditions, MnmC2-dependent activity is required to synthesize mnm^5^s^2^U from the nm^5^s^2^U-tRNA generated by YhcC, most likely due to an absence of an alternative nm^5^(s^2^)U methyltransferase (**Fig. 2B**). This conclusion is consistent with previous studies suggesting that the MnmC1 and MnmC2 domains act independently of each other (14, 35).

It is interesting that *E. coli* relies on YhcC for cmnm^5^s^2^U demodification when MnmC1 is inactive instead of utilizing the “ammonium route” activity of MnmEG, which produces nm^5^s^2^U directly. The dual MnmEG activities have been detected *in vitro*, and previously, it was reported that *E. coli* utilizes the ammonium route during stationary-phase, and the glycine route during exponential growth (14). Indeed, stationary phase seems to be the sole growth condition in which the ammonium route has been reported to be active *in vivo* for any bacterial species. In the case of *E. coli*, it is possible that high levels of glycine in LB medium favor the glycine route over the ammonium route. Previously it was found that *E. coli ΔmnmC* produces cmnm^5^s^2^U but not mnm^5^s^2^U or nm^5^s^2^U in minimal medium containing free ammonia; however yeast extract containing glycine was also included in this experiment (14). In other cases, the ability of MnmEG to directly produce nm^5^s^2^U has been supported through indirect evidence. For instance, there are some bacteria that produce mnm^5^(s^2^)U despite lacking an enzyme capable of cmnm^5^(s^2^)U demodification (MnmC, standalone MnmC1 or MnmL). In other cases, such as in *S. mutans* or *S. pneumoniae*, deleting the gene encoding MnmL does not affect mnm^5^s^2^U production (19). While our genetic experiments were able to replicate nm^5^s^2^U production in stationary phase, we show that, in *E. coli*, this is, in fact due to YhcC activity, and not MnmEG directly producing this intermediate. Therefore, direct evidence for the production of nm^5^s^2^U by MnmEG *in vivo* is lacking in this model organism despite *in vitro* evidence of this activity.

*E. coli* and other bacteria routinely encode parallel enzymes that carry out identical transformations under aerobic versus anaerobic conditions. rSAM enzymes in particular have been observed to be favored by anaerobes and facultative anaerobes to catalyze reactions otherwise catalyzed by oxygen-dependent enzymes in tetrapyrrole synthesis (39). For example, *E. coli* encodes both the oxygen-dependent oxidase HemF and the oxygen-independent dehydrogenase rSAM enzyme HemN, which each catalyze the conversion of coproporphyrinogen III to protoporphyrinogen IX in heme biosynthesis (40, 41). Another key example is the ribonucleotide reductases (RNRs), which catalyze the formation of deoxynucleoside triphosphates for DNA synthesis, where class Ia tyrosine-cysteine radical RNRs utilize O_2_ to generate a reactive complex under aerobic conditions, but class III glycyl radical RNRs (encoded by *nrdD* in *E. coli*) use an O_2_-sensitive [4Fe-4S] cluster and rSAM chemistry to generate a reactive complex under anaerobic conditions (42). Like *yhcC*, *nrdD* transcription is activated by FNR under anaerobic growth conditions (43). Additionally, parallel tRNA hydroxylation pathways and enzymes that utilize divergent O_2_-dependent or independent chemistry ensure efficient biosynthesis of the 5-hydroxyuridine modification and its derivatives in *E. coli* under both aerobic and anaerobic metabolism (44). The work presented here on *yhcC* provides a new example of a dual pathway that allows *E. coli* to efficiently install mnm^5^s^2^U wobble base modifications that promote accurate translation under both aerobic and anaerobic growth conditions. Such parallel pathways are key adaptations that enable bacteria to catalyze essential biochemical transformations under dramatically different environmental and growth conditions by leveraging divergent enzyme chemistries. A phylogenetic analysis further shows that encoding both MnmC and MnmL is common in facultative bacteria, providing evidence that the dual use of these enzymes is a prevalent strategy in this group. However, this seems to be mostly confined to Pseudomonadota bacteria, and the more common strategy for cmnm^5^s^2^U demodification utilized by bacteria is the sole use of either MnmL or MnmC.

## Materials and Methods

### Bioinformatics

The structure of *E. coli* YhcC and *B. subtilis* MnmL proteins was modeled using the AlphaFold3 web server (alphafoldserver.com) and predicted structures were visualized in ChimeraX. To construct the model that included the [4Fe-4S] cluster, 5’dAdo, and methionine, EcYhcC was aligned to *Homo sapiens* Elp3 solved with these cofactors (PDB: 8PTY; ref 33). Multiple sequence alignments were constructed using MAFFT under default settings and visualized in UGENE (46, 47). The large multiple sequence alignment used to detect highly conserved residues was generated from the sequences belonging to the repnode 55 sequence similarity network constructed as described previously, and proteins smaller than 225 and larger than 400 amino acids were discarded (19). Gene neighborhood analysis of YhcC and homologs from related bacteria (Gamma and Betaproteobacteria) was conducted using *fast.genomics* and gene neighborhoods were visualized using Gene Graphics (48, 49).

### Bacterial strain construction and growth conditions

A list of all strains used in this study can be found in Table S1. To ensure all mutants were in an isogenic background, *E. coli* knockout strains were constructed by P1 transduction into *Escherichia coli* str. K12 substr. BW25113 using corresponding mutants obtained from the Keio collection (50). To generate the double *ΔyhcC ΔmnmC::kan* knockout, the kanamycin resistance cassette was first evicted from BW25113 *ΔyhcC::kan* using the FLP recombinase expression plasmid pCP20 (51). For the complementation experiment, the *E. coli yhcC* gene was cloned into pBAD24 by Gibson assembly (New England Biolabs, HiFi assembly kit) utilizing the NcoI restriction site. The pBAD24 vectors containing *mnmLM* and *mnmL* from *Bacillus subtilis* were constructed previously by the Dos Santos lab (19). Site-directed mutants were constructed using the KLD kit (New England Biolabs). Gene expression was induced by the addition of arabinose to 0.02% at the time of inoculation. All strains were grown in LB medium (10 g L^-1^ tryptone, 5 g L^-1^ yeast extract, 10 g L^-1^ NaCl) under either aerobic or anaerobic conditions at 37 °C. Under aerobic conditions, the strains were grown in 50 mL LB from overnight cultures with shaking in 500 mL Erlenmeyer flasks until either mid-log phase (OD_600_ ∼0.5-0.7), late log phase (OD_600_ ∼1.0), or stationary phase (OD_600_ ∼3.0). A Bactron anaerobic chamber (Sheldon Manufacturing) containing a 37 °C incubator in an atmosphere of 90% N_2_, 5% H_2_, 5% CO_2_ was used for anaerobic growth. Plasticware and liquids were brought into the box two days before each growth experiment, and liquids were stirred overnight to allow for gas exchange. Strains were grown on solid LB medium aerobically, then brought into the anaerobic chamber to start the standing overnight cultures. The overnight cultures were subsequently subcultured into 50 mL of LB in Falcon tubes. Kanamycin sulfate (10 μg/mL) was added to agar and overnight cultures when growing the knockout strains, but omitted in the subcultures, and ampicillin (100 μg/mL) was added to select for pBAD24 derived plasmids.

### Cloning and overexpression of YhcC

The *E. coli* YhcC gene was obtained as a codon-optimized gBlock from Integrated DNA Technologies and cloned into a pET28a bacterial expression vector with an N-terminal 6xHis affinity tag and TEV cleavage site using Gibson assembly (New England Biolabs’ HiFi DNA Assembly). The final vector sequence was confirmed using whole plasmid DNA sequencing (Plasmidsaurus). The YhcC pET28a and a pDB1282 plasmid containing the *isc* operon were co-transformed into *E. coli* BL21 (DE3) cells (52, 53). Bacteria were grown in LB medium at 37 °C with shaking until reaching an OD600 between 0.6-0.8 before inducing YhcC expression with 10 μM IPTG. Induced cultures were grown at 18 °C overnight with shaking until cells were harvested at 6,000 x *g* for 40 min at 4 °C, flash frozen with liquid nitrogen, and stored at -80 °C.

### Purification of YhcC

Purification of WT YhcC was carried out in an anaerobic chamber (MBraun) at an oxygen concentration <1 ppm, while YhcC used for binding studies was purified aerobically. Buffers used for purification are as follows: lysis/wash buffer: 50 mM Bis Tris, 500 mM KCl, 2 mM imidazole, 10 mM BME, pH 7.0; wash buffer: 50 mM Bis Tris, 300 mM KCl, 10 mM imidazole, 10 mM BME, pH 7.0; elution buffer: 50 mM, 300 mM KCl, 250 mM imidazole, 10% glycerol, 10 mM BME, pH 7.0; sizing buffer: 50 mM Bis Tris, 500 mM KCl, 5 mM dithiothreitol, pH 7.0. For buffers with 2-mercaptoethanol (BME), 10 mM BME was added immediately before use. All buffers were sparged with argon for 1 hour, moved into the glovebox, and stirred overnight in the anaerobic chamber. YhcC cell pellet was thawed in the glovebox, resuspended in 35 mL of lysis/wash buffer, then lysed in an air-tight sonicator adaptor on ice (Branson Sonifier 450, 4 x 2 min cycles at setting 6, 50% duty cycle). After sonication, lysate was transferred to a centrifuge tube then centrifuged for 45 min at 14,500 rpm (∼24,450 x *g*) and 4 °C. Centrifuge tubes were then brought back into the anaerobic chamber and mixed with 5 mL Ni-NTA resin for 1 hour; Ni-NTA resin was pre-equilibrated with lysis/wash buffer and sparged with argon gas. The Ni-NTA slurry was added into a gravity column and washed with 50 mL of lysis/wash buffer, 50 mL wash 2 buffer, and 50 mL wash 3 buffer. His-YhcC was eluted with 15 mL elution buffer and collected as 1 mL fractions. YhcC was identified by gel electrophoresis (10% SDS-PAGE gel), and YhcC-containing fractions were pooled and concentrated to 2.5 mL using Amicon Ultra 0.5 mL spin columns (10 kDa cutoff) and then purified with size exclusion chromatography on an AKTA Pure 25M with HiLoad 16/600 Superdex 75pg column, pre-equilibrated with 3 CV of anaerobic sizing buffer. SEC fractions were collected inside the anaerobic chamber.

### Chemical Reconstitution of YhcC

SDS-PAGE was used to determine the sizing fractions containing YhcC. Fractions were concentrated and buffer exchanged into reconstitution buffer (50 mM Bis Tris, 300 mM KCl, 10% glycerol, pH 7.0). The protein was diluted to 50 μM with 1 M DTT to reach a final concentration of 5 mM DTT and allowed to incubate at 4 °C for 1 hour. Iron (III) chloride hexahydrate and sodium sulfide nonahydrate were used as Fe and S salts, respectively. A total of 10 equivalents of Fe salt was added in 6 separate aliquots with 5 min between additions. After the final addition of Fe salt, protein rested for 10 min before 10 equivalents of S^2-^ was added in 6 separate aliquots with 15 min between additions. Reconstitution was allowed to continue for 2 hours before being centrifuged for 10 min at 14,000 rpm and the supernatant was buffer exchanged into fresh reconstitution buffer using Amicon Ultra 0.5 mL spin concentrators (10 kDa cutoff) to remove excess Fe/S salts. Presence of the Fe-S cluster was verified by UV-Vis absorbance at 420 nm (**Fig. S5B**). The yellow-brown supernatant was concentrated, flash frozen, and stored at -70 °C.

### Purification of tRNA

tRNA was purified using the phenol-chloroform method utilizing TRIzol (Invitrogen) and the NucleoBond RNA/DNA 80 kit (Macherey-Nagel). Cultures reaching the desired growth phase were immediately placed on ice and centrifuged at 3,700 x *g* at 4 °C. Cell pellets were then resuspended in 1 mL TRIzol and placed on a rocker for 15 minutes. 200 μL chloroform was added, and the cell extracts were then centrifuged at 15,000 x *g* for 15 minutes to separate the organic and aqueous phases. Aqueous phases were transferred to new tubes and to these, one volume of cold isopropanol was added. The samples were then placed at -20 °C for 30-60 minutes or 4 °C overnight. Samples were centrifuged at 15,000 x *g* for 20 minutes and the supernatants were discarded. In the NucleoBond purification, the RNA pellets were resuspended in 6 mL of buffer R0 (100 mM Tris/acetate, 15% ethanol, pH = 6.3) and applied to Nucleobond columns. The tRNA was then purified according to the manual, except tRNA was eluted with 1.1 mL buffer R2 (100 mM Tris/acetate, 15% ethanol, 900 mM KCl, pH = 6.3) diluted to 0.75 M KCl with buffer R0. The purified tRNA was precipitated by addition of 916 μL cold isopropanol followed by incubating at 4 °C overnight. Precipitated tRNA was centrifuged at 15,000 x *g* for 20 minutes, and the pellets were washed twice with 80% ethanol, dried, and resuspended in pure water.

To prepare cmnm^5^s^2^U-enriched tRNA for the YhcC and MnmC1 activity assays describe below, tRNA was purified from *E. coli* T7 Express *ΔmnmC* + pSGAT2-tRNA^Glu^-II strain, constructed previously by the Carell lab, which provided the strain to us (17). The strain was grown aerobically with shaking in 250 mL LB medium to an OD_600_ of 1.34, and tRNAs were purified as described above, except large RNAs were precipitated by the addition of NaCl to 1 M prior to the NucleoBond purification step.

### Quantification of tRNA modified nucleosides

Relative levels of nucleoside modifications from tRNA purified from *E. coli* cultures were analyzed by chromatography-coupled mass spectrometry methods (LC-MS) as previously described (19). For each analysis, 40 μg of denatured tRNA were digested with nuclease P1 and alkaline phosphatase prior to LC-MS detection. Dihydrouridine standard (Dalton) was run as a control for normalizing data as it is present in lower abundance, similar to the modifications analyzed in this study. The relative levels of each modification were determined by the area of mass abundance of each modification by mass abundance of dihydrouridine in the same sample. The desired analytes D ([M+H]^+^:247.0925), nm^5^s^2^U ([M+H]^+^: 290.0805), cmnm^5^s^2^U ([M+H]^+^: 348.086), and mnm^5^s^2^U ([M+H]^+^:304.096) were determined within 5 ppm accuracy.

### *In vitro* transcription of unmodified *E. coli* tRNA

The *E. coli* tRNA^Lys^_UUU_ sequence (5’-GGGTCGTTAGCTCAGTTGGTAGAGCAGTTGACTTTTAATCAATTGGTCGCAGGTTCGAATCCTGCACGACCCACCA-3’) was obtained from the tRNADB-CE tRNA sequence database (https://trna.ie.niigata-u.ac.jp/cgi-bin/trnadb/whole_detail.cgi?SID=21833864&). dsDNA template with added 5’ T7 promoter sequence for this tRNA sequence was obtained from Integrated DNA Technologies; template dsDNA sequence = 5’-**TAATACGACTCATATA**GGGTCGTTAGCTCAGTTGGTAGAGCAGTTGACTTTTAATC AATTGGTCGCAGGTTCGAATCCTGCACGACCCACCA-3’). The tRNA sequence was *in vitro* transcribed using T7 polymerase in a 500 μL reaction: 1 μM DNA template, 5 mM rNTPs, 5 mM DTT, 0.05% v/v Triton X-100, 6 U/mL thermostable inorganic pyrophosphatase, and 0.15 mg/mL T7 RNA polymerase in 1 X T7 buffer (New England Biolabs). The reaction was incubated overnight at 37 °C before DNA template was digested with TURBO DNase (Thermo Fisher Scientific) for 1 hour at 37 °C. Subsequently, tRNA was purified using denaturing urea PAGE: tRNA product band was gel extracted (RNA extraction buffer: 0.6 M NaOAC, 1 mM EDTA, pH 6.0, 0.01% SDS) and precipitated with 1/10 volume 3 M NaOAc and 3x sample volume 100% ethanol at -20 °C. Precipitated tRNA was pelleted by centrifugation, redissolved in water, and annealed (heated at 80 °C for 2 min, 60 °C for 2 min, then 100 mM MgCl_2_ was added to a final concentration of 10 mM, and cooled on ice for 30 min). Annealed tRNA was aliquoted, flash frozen with liquid nitrogen, and stored at -80 °C until ready for use.

### Electrophoretic Mobility Shift Assays (EMSAs)

Triplicate YhcC-tRNA binding reactions were prepared with an unmodified tRNA^Lys^_UUU_ concentration of 200 nM and YhcC concentrations ranging from 50 – 0.26 μM protein (1.5X dilution series) in YhcC reconstitution buffer containing 10% glycerol (50 mM Bis Tris, 300 mM KCl, 10% glycerol, pH 7.0). Binding reactions were prepared by combining 2X protein with 2X tRNA to a final volume of 10 μL, incubated at 4 °C for 1.5 hours, and then 7 μL was loaded into each well of a 5 % MP TBE gel (Bio-Rad). EMSA gels were run in 0.5X TBE at 4 °C and 8 mA. Staining of tRNA was carried out with SYBR Gold in 0.5X TBE buffer for 30 min with gentle shaking. Gels were imaged on a Fluorochem Q imager (EtBr setting). Gel images were then analyzed using ImageJ to quantify bound and unbound tRNA species. Binding curves were fit to a Hill coefficient plot using Prism.

### MnmC1 tRNA modification assay

Purified MnmC1 was obtained from the Carell lab and MnmC1 activity assays were carried out as previously described (17). but under aerobic (benchtop) or anaerobic (inside a glovebox with degassed buffers) conditions. Equal volumes of 2X tRNA purified from *E. coli* Δ*mnmC* cells as described above in water (7 μM) and 2X MnmC1 enzyme (0.1 μM) in 2X reaction buffer (120 mM Tris-HCl, pH 8.0, 40 mM NH_4_Cl, 5 mM KCl, and 1.3 mM MgCl_2_) were mixed at 37 °C to initiate the reaction. Final tRNA and enzyme concentrations in the reaction were 3.5 μM and 0.05 μM, respectively, in 1X reaction buffer (60 mM Tris-HCl, pH 8.0, 20 mM NH_4_Cl, 2.5 mM KCl, and 0.65 mM MgCl_2_). Reaction timepoints were quenched by adding methanol to the reaction aliquot at a 1:1 volume ratio. Quenched reaction samples were cleaned up using the Zymo Research Clean and Concentrate RNA kit, eluting isolated tRNA in 15 μL water prior to using an NEB nucleoside digestion mix (0.5 μL enzyme with 1.72 μL 10X digestion buffer). Digestion occurred overnight at 37 °C prior to being transferred to the UHPLC-MS block for analysis by Agilent Bio-Inert 1260 Infinity II UHPLC system with Infinity Lab LC/MSD. The UHPLC was equipped with the Agilent Zorbax SB-Aq Rapid Resolution HD using a mobile phase with 0.1% formic acid in water (A) or 0.1% formic acid in acetonitrile (B), and detected using positive ionization mode. The gradient was as follows for detection of cmnm^5^s^2^U (m/z = 348.1, 1.1 min) and nm^5^s^2^U (m/z = 290.1, 0.607 min) at 0.4 mL/min: 0-2.50 min, 100% A; 2.50-6.00 min, 20% A/80%B; 6.00-10.5 min, 100% A. Concentrations of substrate cmnm^5^s^2^U and product nm^5^s^2^U nucleosides in quenched reaction samples were calculated relative to calibration curves made with synthetic nucleoside standards, which were synthesized as previously described (54).

### YhcC tRNA modification assay

YhcC activity assays were carried out under anaerobic conditions (<1 ppm oxygen) with final reaction mix concentrations of: 1 μM reconstituted YhcC, 25 μM *S*-adenosyl methionine, 0.5 mM dithionite, and 3.5 μM tRNA purified from Δ*mnmC E. coli* cells, as described above for MnmC1 assays. A 2X solution of YhcC (2 μM) and cofactors (50 μM S-adenosyl methionine, 1 mM dithionite) in 2X reaction buffer (120 mM Tris-HCl, pH 8.0, 40 mM NH_4_Cl, 5 mM KCl, and 1.3 mM MgCl_2_) was incubated at 37 °C for 10 min and reactions were initiated by 1:1 addition of a 2X solution of tRNA (7 μM) in water. All activity assays were run in triplicate with time points collected by quenching 15 μL of reaction mix with 1:1 methanol. Time points were collected at 0, 90, 300, 600, 1800, 5400 s. Quenched samples were removed from the anaerobic chamber and cleaned up using Zymo Research Clean and Concentrate RNA kit, eluting isolated tRNA in 15 μL water. NEB nucleoside digestion mix (0.5 μL enzyme mix with 1.72 μL 10X digestion buffer) was added and allowed to digest overnight at 37 °C. Reactions were transferred to UHPLC-MS block for analysis by Agilent Infinity series and concentrations of substrate cmnm^5^s^2^U and product nm^5^s^2^U nucleosides were calculated as detailed above for MnmC1 assays.

### Analysis of MnmL and MnmC occurrence and association with oxygen requirement

COG and KO incidence per orthologous family were retrieved (accessed July 2026) (31, 32). The organisms shared between the KEGG KO and COG databases were identified and used to define a benchmark set of 1,322 bacterial genomes. BactoTrait database (accessed June 2026) and direct BacDive (accessed June 2026) were used to compile physiological trait data for these benchmark genomes (33, 34). Physiological oxygen requirement data was converted from discrete categories into numeric representations appropriate for heatmap visualization. Oxygen requirement categories and their corresponding encodings are as follows: obligate anaerobe = 0, anaerobe = 1, facultative anaerobe = 2, facultative aerobe = 3, aerobe = 4, obligate aerobe = 5. The terms “oxygen requirement” and “oxygen consensus” are used to refer to the biological category and the numeric representations of those categories, respectively. To improve trait completeness while preserving prior curation, existing accession-based consensus calls were retained, and only missing consensus slots were filled from legacy curation records (unpublished; manually curated from JGI-IMG, BacDive, and individual published organism characterizations; accessed 2023) mapped by unique NCBI Tax ID at the assembly level (55).

## Acknowledgements

This work was funded by the National Institutes of Health (Grants R35GM156215 to VdC-L and R35GM143000 to JSM, which funded key instrumentation used in this study). The content is solely the responsibility of the authors and does not necessarily represent the official views of the National Institutes of Health. This material is based upon work supported by the National Science Foundation under Grant No. 2339759 to JSM and Grant No. MCB-1716535 to PDS; preliminary data collection was also funded by a University of Delaware General University Research (GUR) award to JSM.

## Supplemental Tables

**Table S1.** List of strains used in this study.

| Strain | Genotype | Reference |
| --- | --- | --- |
| KSB1005 | BW25113 $\Delta mnmC::kan$ | This work |
| KSB1007 | BW25113 $\Delta yhcC::kan$ | This work |
| KSB1019 | BW25113 $\Delta yhcC \Delta mnmC::kan$ | This work |
| KSB1025 | BW25113 $\Delta yhcC::kan$ + pBAD24 | This work |
| KSB1029 | BW25113 $\Delta yhcC::kan$ + pBAD24 *EcyhcC* | This work |
| KSB1093 | BW25113 $\Delta yhcC::kan$ + pBAD24 *BsmnmL* | This work |
| KSB1095 | BW25113 $\Delta yhcC::kan$ + pBAD24 *EcyhcC_C33A* | This work |

**Table S2.**
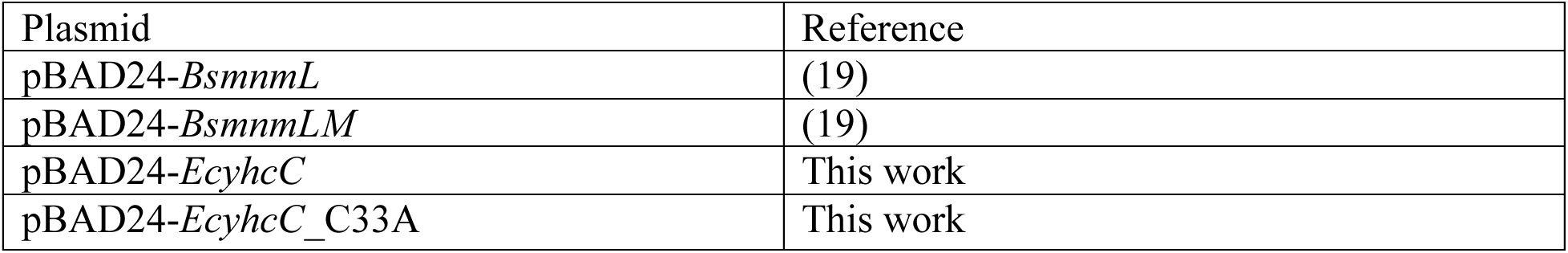
List of plasmids used in the study.

| Plasmid | Reference |
| --- | --- |
| pBAD24-*BsmnmL* | (19) |
| pBAD24-*BsmnmLM* | (19) |
| pBAD24-*EcyhcC* | This work |
| pBAD24-*EcyhcC_C33A* | This work |

**Table S3.**
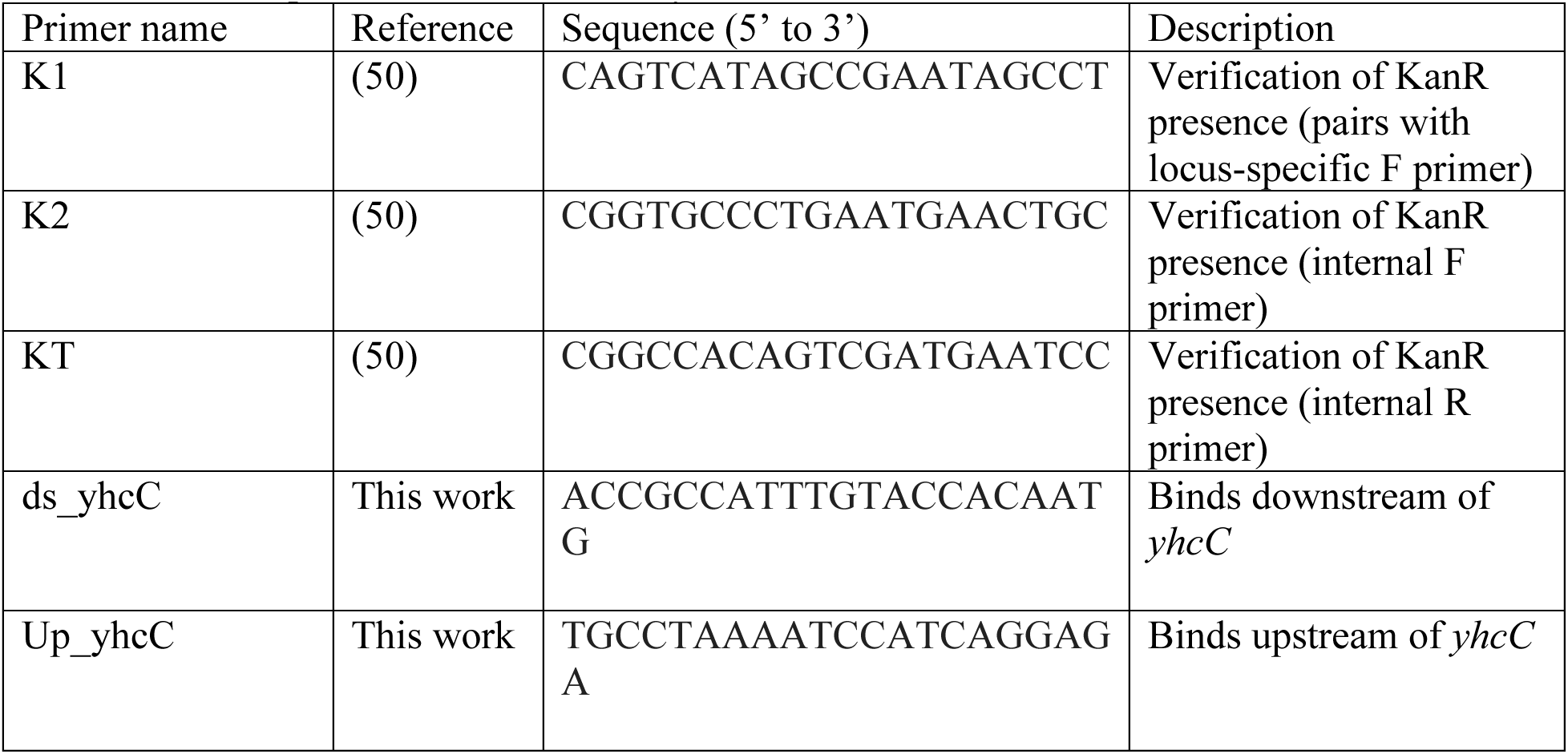
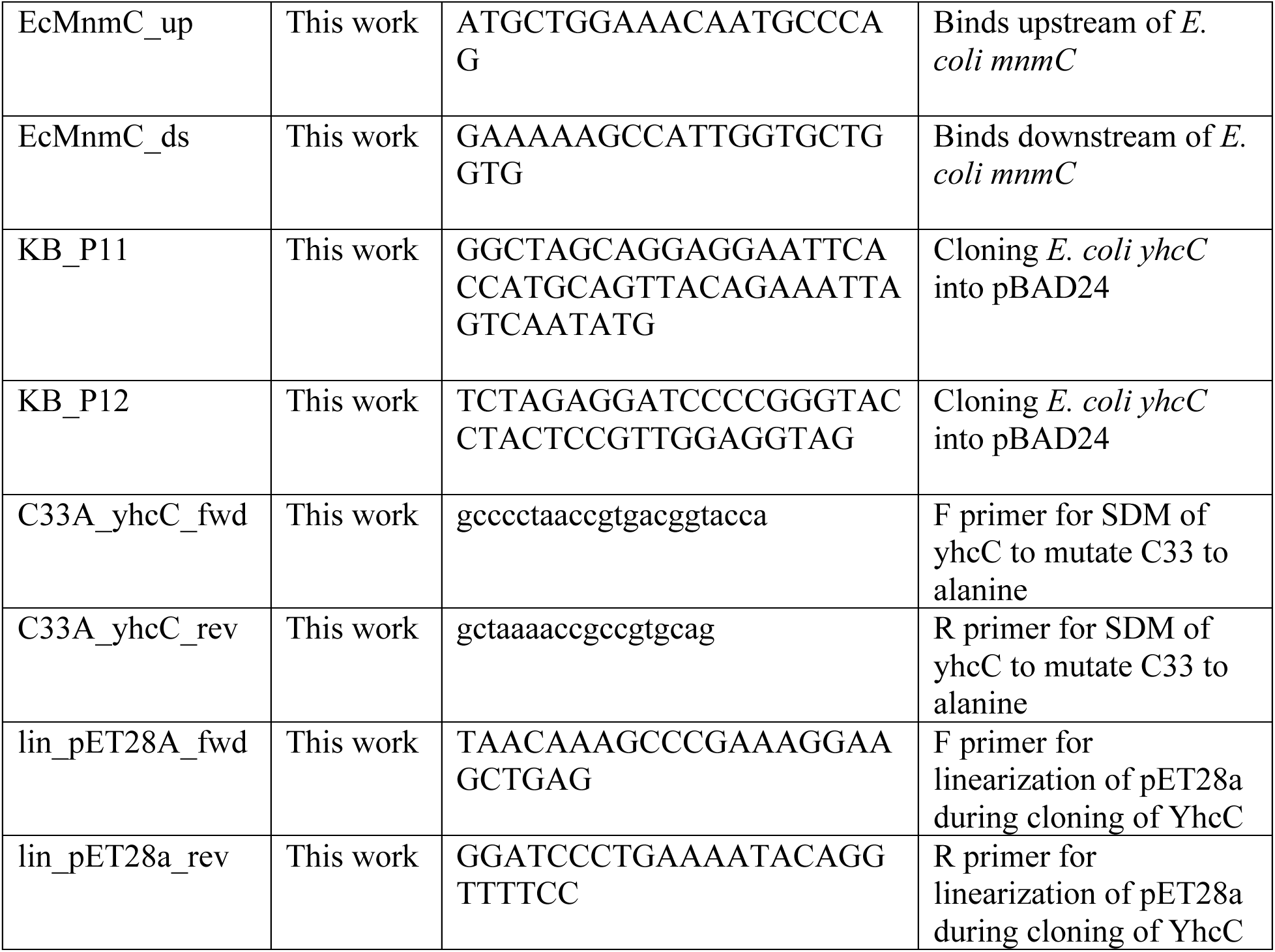
List of primers used in the study.

## Supplemental figures

**Figure S1.**
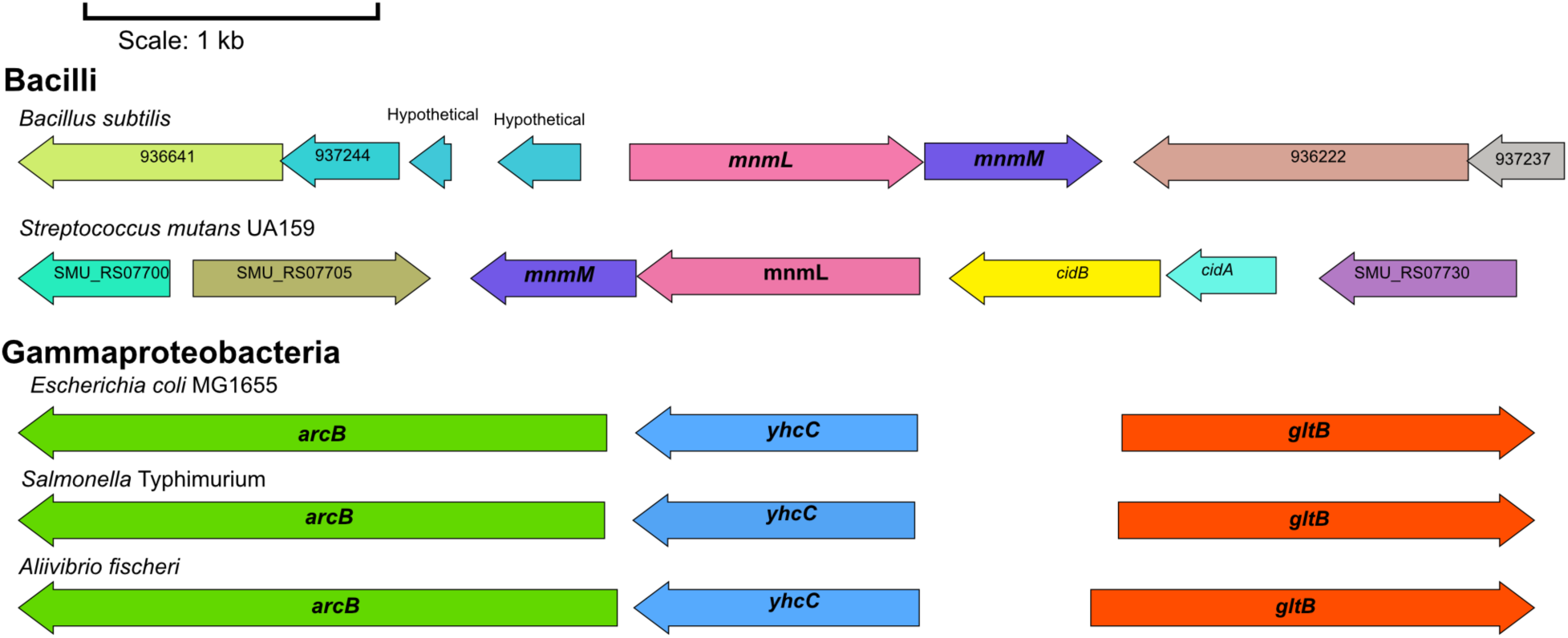
As previously reported, *mnmL* genes strongly associate with *mnmM* in Gram-positive bacteria (Bacilli, top) (19). In contrast, *yhcC* homologs from *E. coli*and related Gammaproteobacteria physically cluster next to the *arcB* redox sensor (Gammaproteobacteria, bottom).

**Figure S2.**
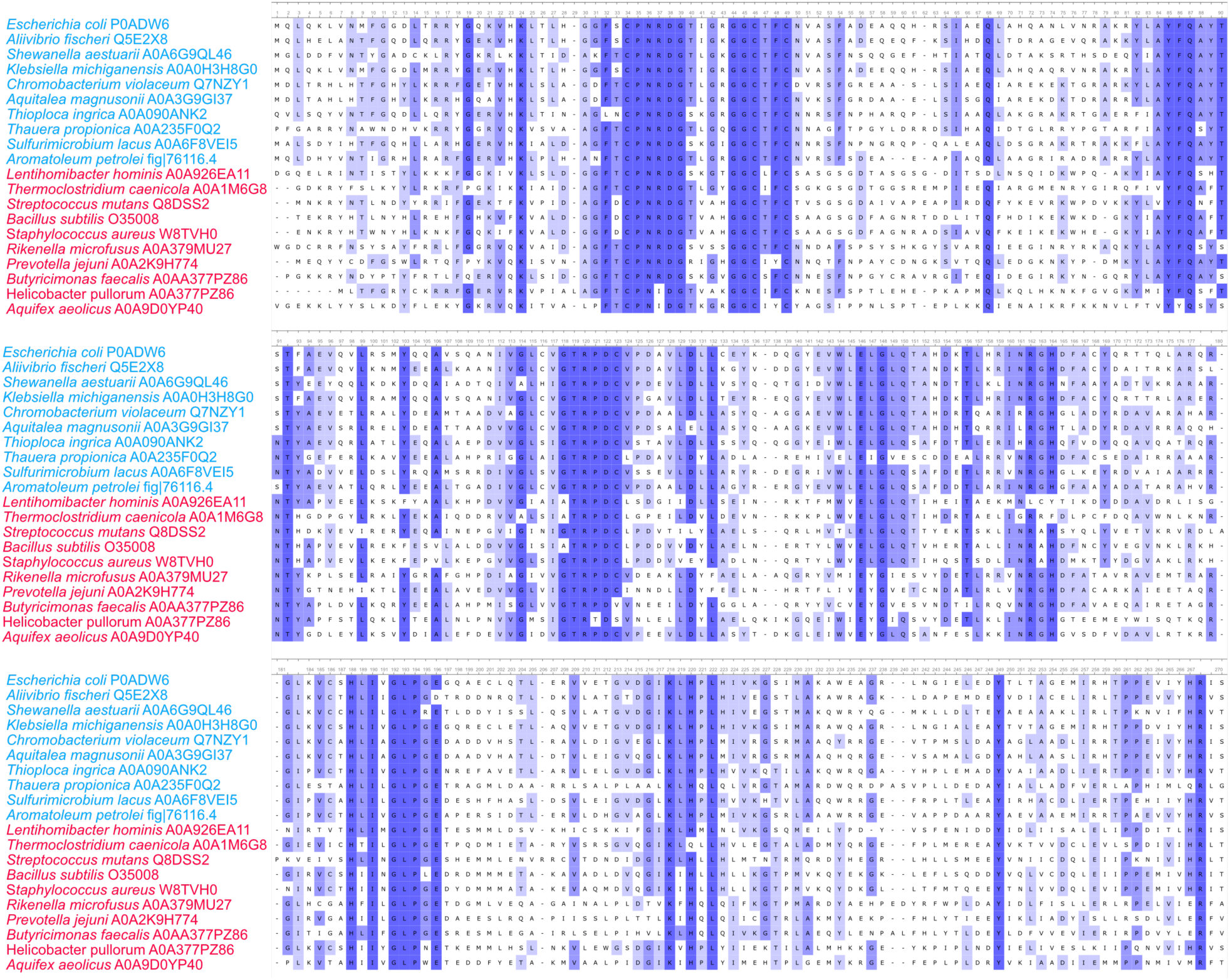
A multiple sequence alignment of YhcC/MnmL proteins from organisms encoding bifunctional MnmC (blue) and organisms that do not encode MnmC (red). There do not appear to be any primary sequence-specific differences between the two sets of proteins. The sequences are colored based on conserved identity.

**Figure S3.**
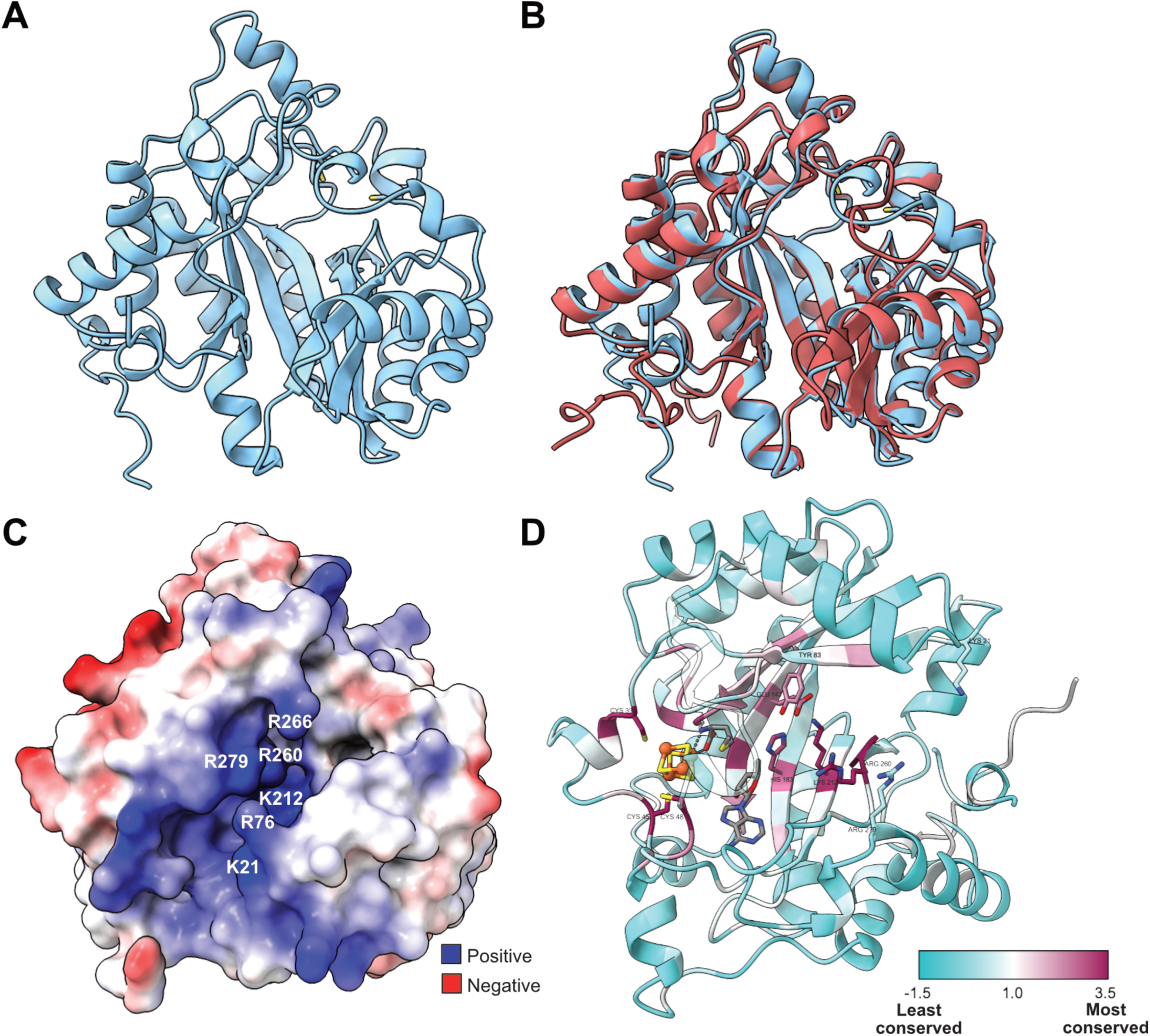
Structural analyses reveal that YhcC from *E. coli* (EcYhcC) and MnmL from *B. subtilis* (BsMnmL) exhibit very similar structures. **A**: Overall predicted structure of EcYhcC showing the rSAM cysteine motif (sulfur atoms in yellow). **B**: Predicted EcYhcC overlaid with the AlphaFold-predicted structure of BsMnmL. **C**: Electrostatic map of EcYhcC shows a positively-charged cleft that leads towards the predicted active site of the enzyme. **D**: Structure highlighting conserved residues found either in the predicted enzyme active site or in the positively-charged cleft. The active site is predicted based on the radical SAM cysteine residues; an [4Fe-4S] cluster, methionine, and 5’-deoxyadenosine were docked into the structure by aligning with human Elp3 (PDB: 8PTY). Note that this structure is rotated relative to **Fig. S3A-C** to better highlight conserved residues.

**Figure S4.**
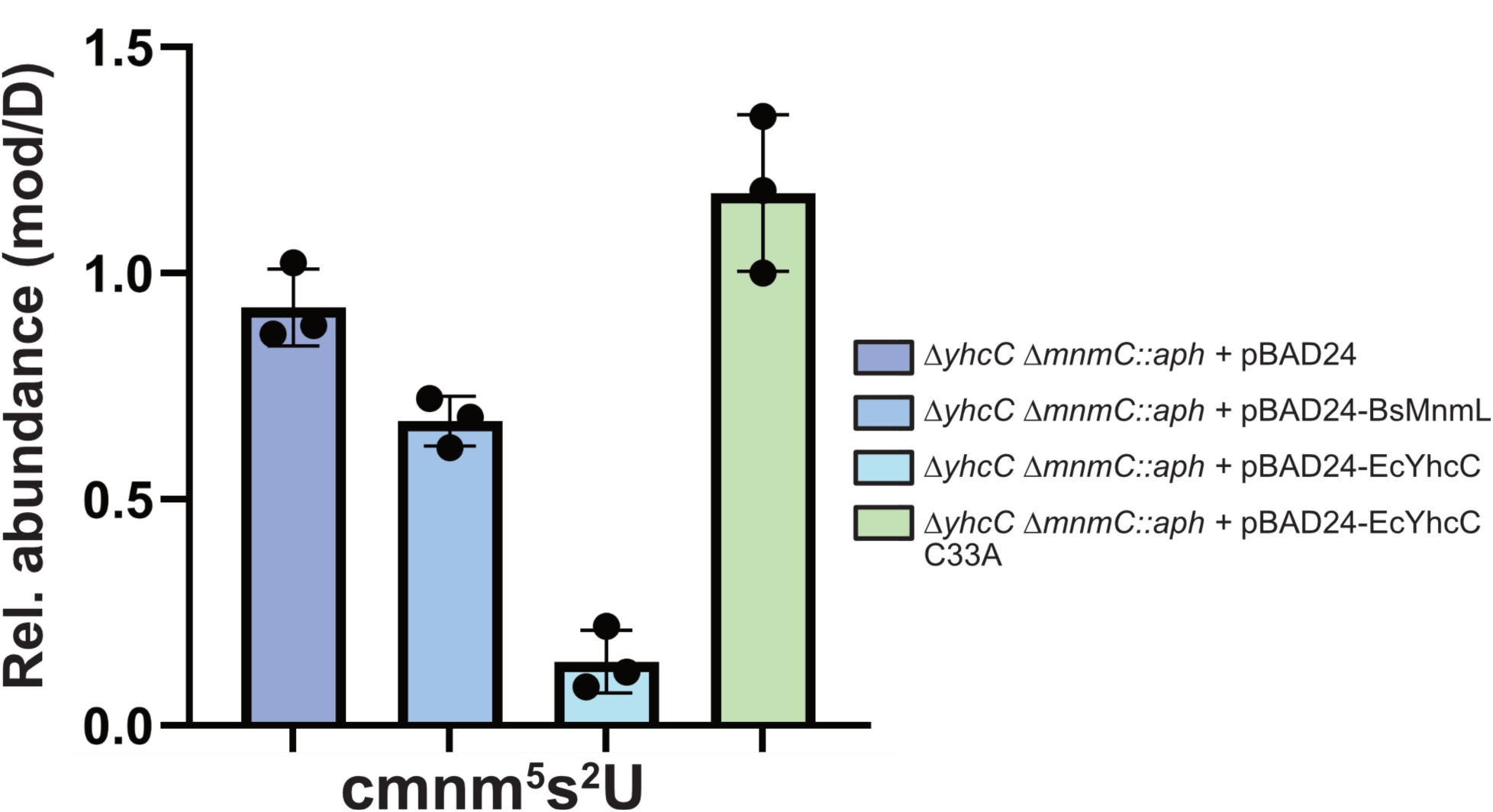
Aerobic complementation of *yhcC* and *mnmC*. The double *ΔyhcC ΔmnmC* strain was transformed with empty pBAD24, or pBAD24 with either *Bacillus subtilis mnmL* (BsMnmL), *E. coli yhcC* (EcYhcC) or *E. coli yhcC* carrying the C33A mutation (EcYhcC C33A). The strains were grown under aerobic conditions to mid-log phase before tRNAs were purified and subsequently analyzed for the presence of cmnm^5^s^2^U, mnm^5^s^2^U, and nm^5^s^2^U, alongside the non-thiolated versions. Only cmnm^5^s^2^U was detected in these strains.

**Figure S5.**
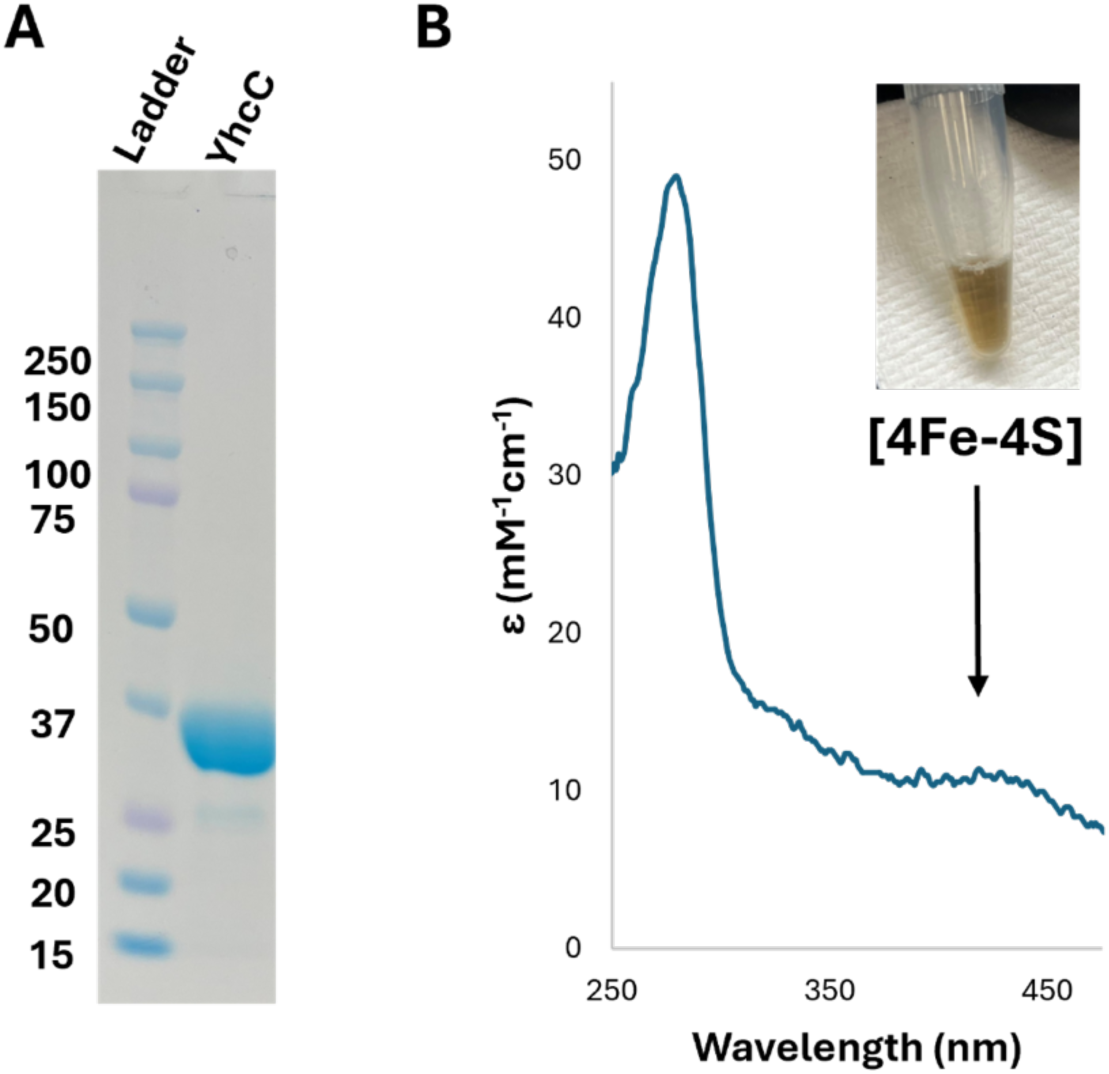
Recombinant His-tagged YhcC was anaerobically purified (**A**) and the [4Fe-4S] cluster was chemically reconstituted *in vitro*. The UV-vis spectrum of purified, reconstituted YhcC enzyme at 5 μM (**B**) shows a characteristic absorption at 420 nm, indicative of [4Fe-4S] formation.

**Figure S6.**
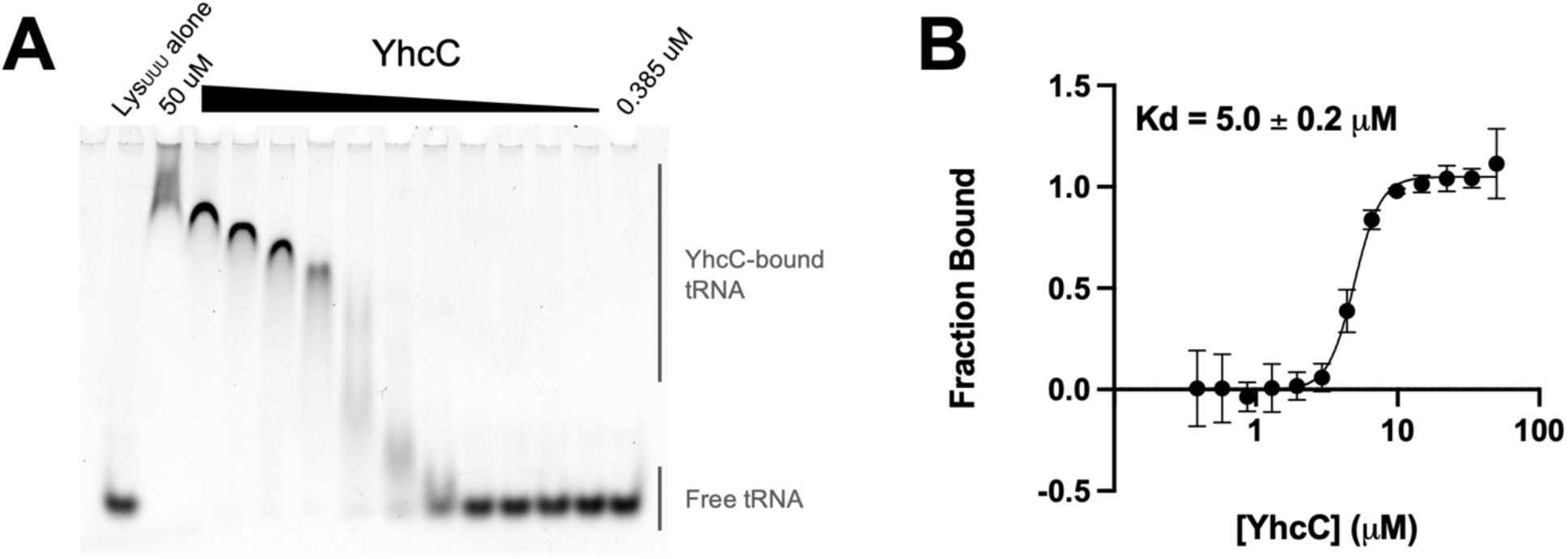
tRNA binds YhcC with low micromolar affinity *in vitro*. **(A)** EMSA gel showing binding of *E. coli* LysUUU tRNA to purified YhcC. **(B)** Quantification of fraction bound versus YhcC concentration from EMSA fits to a *K*_D_ of 5 μM.

**Fig. S7.**
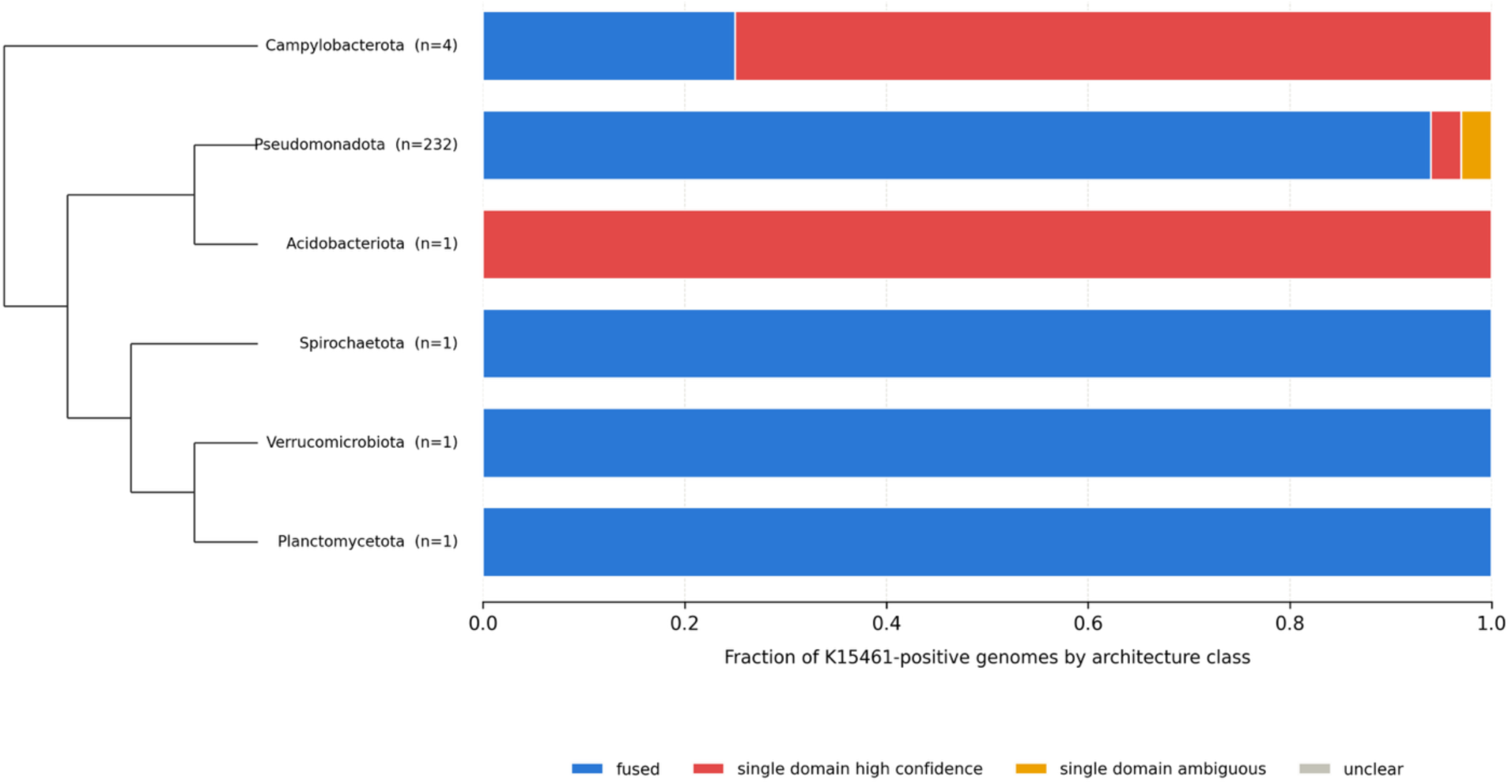
MnmC (K15461) is found in six phyla, however some sequences in this dataset were annotated to contain either only the oxidase domain or only the methyltransferase domain by the Pfam database. These proteins are classified here as either single domain “high confidence” when less than 500 amino acids, or “ambiguous” when more than 500 amino acids in length.

**Figure S8.**
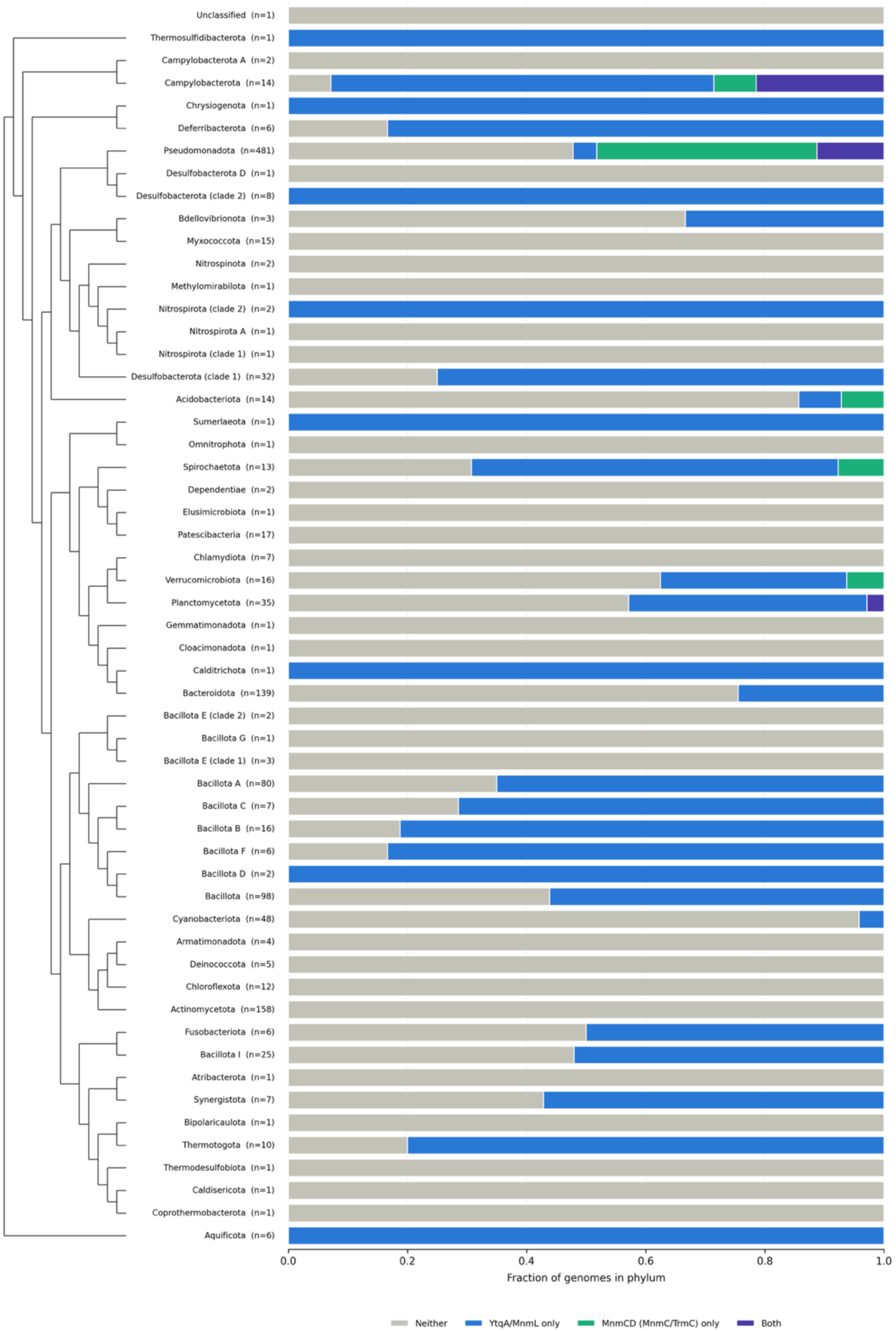
Phylogenetic distribution of MnmL and MnmC in 1,322 representative genomes spanning 54 phylogenetic groups. MnmL is found in 28 groups, whereas MnmC is found in six, although the majority of the latter homologs are restricted to the Pseudomonadota phylum (see **Fig. S7**).

**Figure S9.**
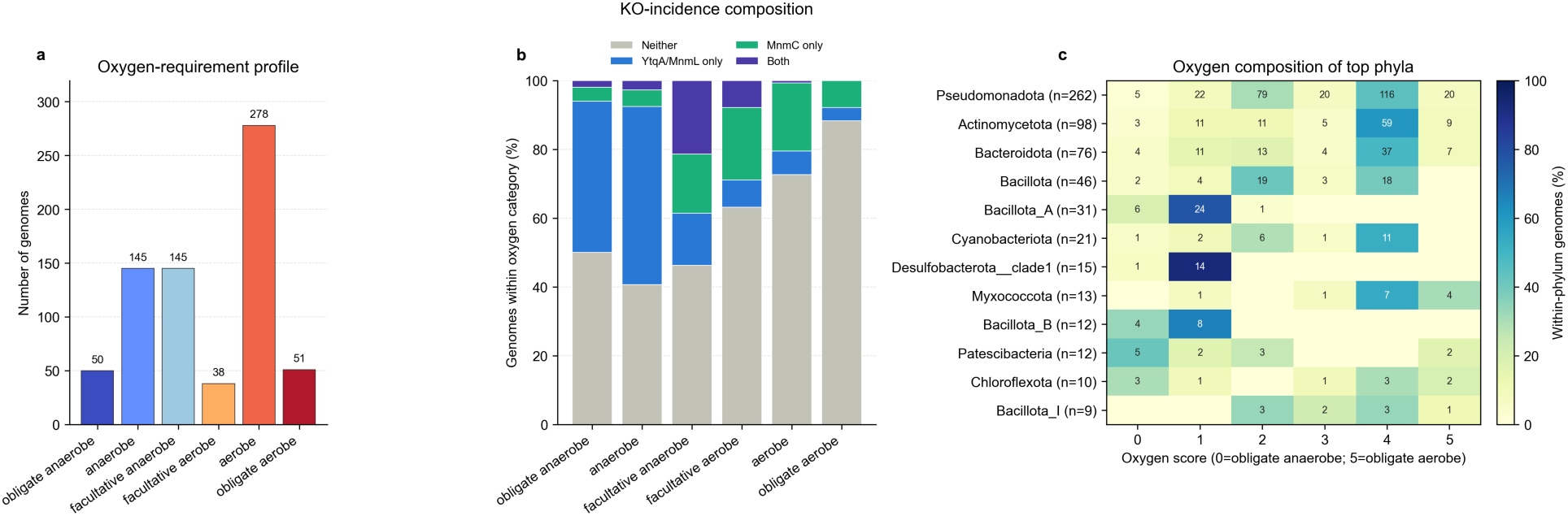
Summary of the 707 representative bacteria containing oxygen utilization data. **A**: The number of bacteria belonging to each of the six oxygen utilization categories. **B**: Incidence of MnmL and MnmC in the 707 organisms by oxygen utilization. **C**: Oxygen utilization breakdown of the 12 most abundant phyla in the dataset.

